# Evolutionary plasticity of regulatory–catalytic coupling in mono- and poly-ADP-ribosyl transferases

**DOI:** 10.64898/2026.10.17.759577

**Authors:** Makwana Milee, Khare Shambhavi, Shanavas P Greeshma, Rahman Ghazala, Bhargava Anamika, Eerappa Rajakumara

**Affiliations:** Macromolecular Structural Biology Lab, Department of Biotechnology, Indian Institute of Technology Hyderabad, India; Sri Aurobindo Institute of Medical Science, India; Cell Signalling and Ion Channel Biology Lab, Department of Biotechnology, Indian Institute of Technology Hyderabad, India

**Author notes:** Corresponding Author, Correspondence to Eerappa Rajakumara. These authors contributed equally (Makwana Milee, Khare Shambhavi).

## Abstract

The evolution of modular proteins enables biological systems to diversify regulatory functions while preserving conserved catalytic activities. ADP-ribosyltransferases (ARTs), including mono-ADP-ribosyltransferases (MARylating enzymes) and poly-ADP-ribosyltransferases (PARylating enzymes), provide an ideal model for investigating how regulatory and catalytic domains co-evolve. Although ART catalytic domains are highly conserved, their regulatory architectures vary extensively, raising the question of whether regulatory domains determine catalytic activation, allosteric communication, and enzymatic output. Here, we systematically examined regulatory–catalytic interoperability by reconstituting evolutionarily distant ARTs from mammals, plants, and bacteria. We show that the regulatory domain of the MARylating enzyme human PARP3 (hPARP3) functionally complements the catalytic domains from the PARylating enzymes human PARP1 (hPARP1), human PARP2 (hPARP2), and Arabidopsis PARP2 (atPARP2), restoring robust DNA-break-dependent allosteric coupling for catalytic activation in response to both phosphorylated double-strand and single-strand DNA breaks. Complementation stimulates NAD⁺ consumption, promotes cis- and trans-poly(ADP-ribosyl)ation, and is accompanied by direct physical interactions between heterologous regulatory and catalytic domains. Unexpectedly, the strongest activation was observed between the hPARP3 regulatory domain and the atPARP2 catalytic domain, demonstrating conservation of productive allosteric communication across both enzymatic classes and species boundaries. In contrast, regulatory domains from PARylating ARTs failed to activate the catalytic domain of MARylating hPARP3, revealing a striking asymmetry in regulatory–catalytic compatibility. These findings identify conserved and specialised modes of allosteric communication within the ART superfamily and establish the concepts of general and specific combining ability as a framework for understanding modular protein evolution. More broadly, our results demonstrate that conserved regulatory–catalytic signalling networks can be functionally transferred between divergent enzymes, providing a foundation for mapping allosteric communication across ARTs and for engineering synthetic ADP-ribosylation systems with programmable regulatory inputs and catalytic outputs.

## Introduction

Modular protein architectures are among the most successful evolutionary strategies for generating functional diversity while preserving essential biochemical activities [^1–4^]. Across biological systems, catalytic or functional domains are often highly conserved to maintain core enzymatic functions, whereas regulatory and ligand-binding domains evolve more rapidly to confer context-specific control over activity, substrate recognition, cellular localisation, allosteric regulation, and signalling specificity [^5,6^]. This separation of catalytic function from regulatory control enables organisms to adapt existing molecular frameworks to new cellular contexts, environmental challenges, and developmental demands without altering the underlying catalytic machinery [^5,7,8^]. Consequently, catalytic domains are frequently constrained by strong purifying selection, whereas accessory domains serve as evolutionary platforms for innovation [^9–11^].

Protein kinases provide a classic example of this principle. Although the kinase catalytic core is remarkably conserved throughout eukaryotic evolution, kinases possess highly diverse regulatory modules, including SH2, SH3, PH, PDZ, leucine-rich repeat, and other interaction domains that determine signalling inputs, subcellular localisation, and substrate selection [^12–17^] (Supplementary Fig. 1A). Similarly, receptor tyrosine kinases maintain a conserved intracellular kinase domain while evolving distinct extracellular ligand-binding regions that recognise diverse growth factors and cytokines [^18–21^]. G protein-coupled receptors (GPCRs) illustrate an analogous strategy in which a conserved seven-transmembrane signalling core has diversified through variation in extracellular and intracellular regulatory elements, enabling the detection of stimuli ranging from photons and odorants to neurotransmitters, peptides, and hormones. Such modularity has been a major driver of signalling complexity throughout evolution [^22–25^] (Supplementary Fig. 1B).

The emergence of modular proteins is facilitated by evolutionary processes, including gene duplication, exon shuffling, recombination, and horizontal transfer of domains [^26–28^]. These mechanisms allow pre-existing functional modules to be recombined into novel architectures while retaining catalytic competence [^28,29^]. As a result, modular enzymes and receptors have become important systems for understanding how conserved catalytic cores acquire new biological functions through changes in regulatory domains. Experimental approaches based on cross-complementation and domain swapping have proven particularly powerful for dissecting the contributions of individual modules and for determining how regulatory and catalytic domains communicate to generate specific biological outputs [^29–33^].

Among modular signalling enzymes, ADP-ribosyltransferases (ARTs), commonly referred to as PARPs, represent an outstanding model for investigating the relationship between conserved catalytic mechanisms and regulatory diversification [^34–38^]. ART family proteins possess a conserved catalytic (CAT) domain that transfers ADP-ribose from NAD^+^ onto target proteins, yet exhibit remarkable variation in their regulatory architectures [^34,36–43^]. Different family members contain individual or combinations of zinc-finger domains, WGR domains, BRCT domains, WWE domains, RNA-binding motifs, ankyrin repeats, SAP domains, macrodomains, OB-folds, and other accessory modules [^34,44–46^] (Fig. 1A and Supplementary Fig. 1C). These domains govern DNA damage recognition, substrate recruitment, chromatin interactions, RNA binding, subcellular localisation, and protein-protein interactions, thereby enabling ART family members to participate in diverse biological processes including DNA repair, transcriptional regulation, development, apoptosis, stress responses, telomere maintenance, and innate immunity [^34,47–50^] (Supplementary Fig. 1 C).

**Fig. 1.**
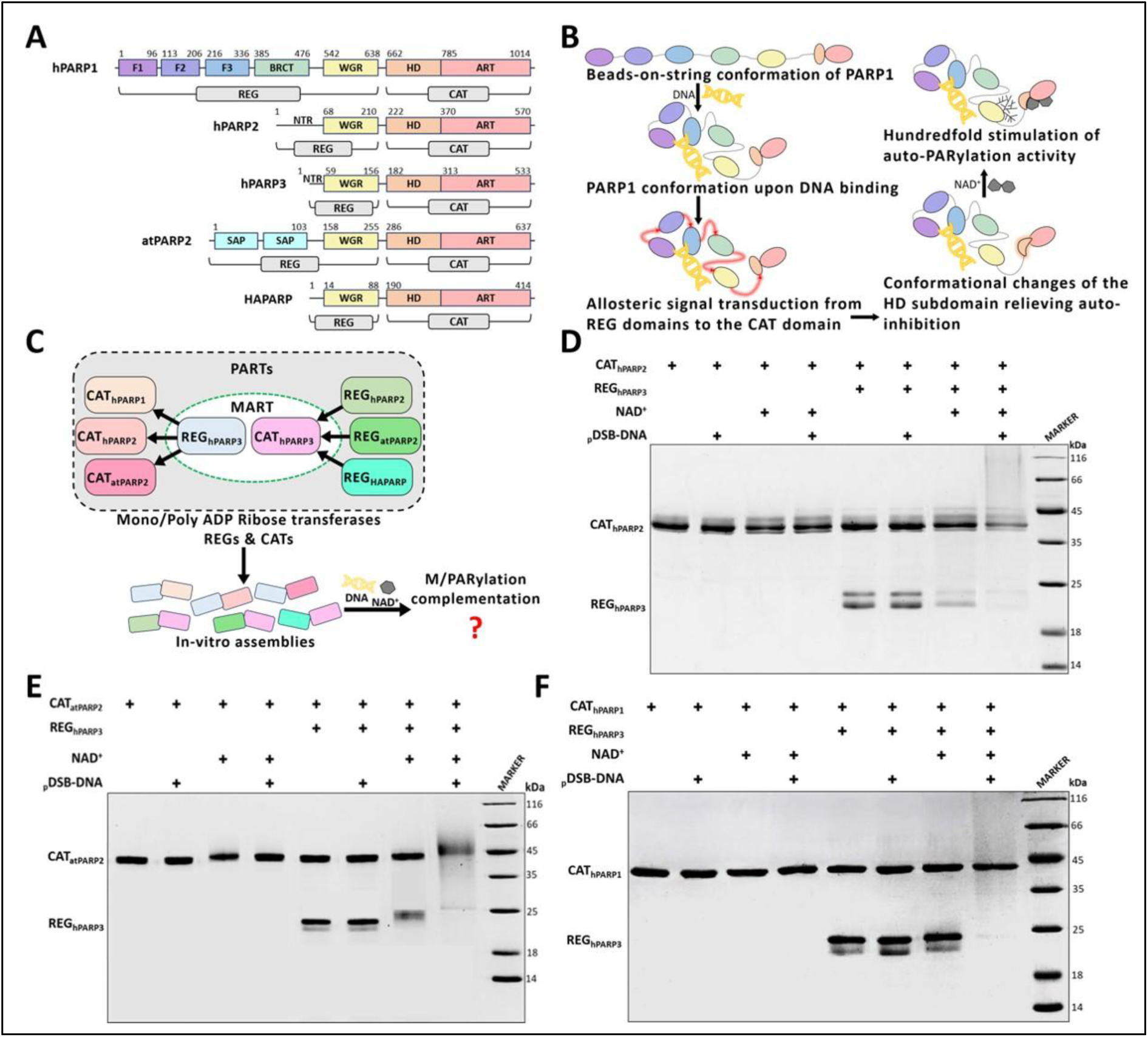
Domain architecture of Poly ADP ribosyltransferases used in the study, experimental summary and cross-complementation PARylation assays. (A) Domain architecture of MART (hPARP3) and PART (hPARP1, hPARP2, atPARP2 and HAPARP) PARPs used in this study. (B) Schematic representation of the relaying of allosteric signal, upon DNA break binding, from DNA breaks recognition domains to the regulatory domains to the ART domain within the CAT domain through the HD subdomain for allosteric activation of catalytic activity of PARP1. Note conformational change in the HD domain upon DNA-dependent assembling of PARP1. Domains of PARP1 are represented in the colour scheme: Purple (Zinc finger domains), Green (BRCT), Yellow (WGR), Peach (HD), Orange (ART) (C). Experimental summary showing different combinations of REG and CAT domains from MART and PARTs of PARPs used in the study with in vitro assemblies to evaluate M/PARylation complementation activity. SDS-PAGE PARylation assay of cross-complementation of (D) CAT_hPARP1_, (E) CAT_hPARP2_, (F) CAT_atPARP2_, in the presence of an allosteric ligand pDSB-DNA, by REG_hPARP3_.

Despite this regulatory diversity, accumulating evidence suggests that several members of the ART family employ related mechanisms of allosteric regulation [^45,51–55^] (Fig. 1B). In the DNA damage-responsive enzymes PARP1, PARP2 and PARP3, recognition of allosteric ligands, DNA breaks, by regulatory domains generates allosteric signals that are transmitted through a regulatory WGR domain to the helical domain (HD), an autoinhibitory and a regulatory subdomain embedded within the catalytic domain [^56,57^] (Fig. 1B). Destabilisation of the HD promotes NAD^+^ access to the active site and stimulates catalytic activity [^57^] (Fig. 1B). Similar regulatory principles appear to operate in response to other allosteric ligands [^36,40–43,45,51,58–60^] including proteins, RNA, poly(ADP-ribose) (PAR), and post-translational modifications, suggesting the existence of a conserved regulatory axis linking diverse sensory domains to a common catalytic output. However, the extent to which regulatory domains dictate catalytic activation, allosteric signal propagation, and enzymatic output across the broader ART superfamily remains unknown.

This question is particularly intriguing because mono-ADP-ribosylating and poly-ADP-ribosylating transferases (MARTs and PARTs) share a highly conserved catalytic fold yet generate fundamentally different biochemical products (Fig. 1A [^34,56,61,62^]. Mono-ADP-ribosyltransferases such as human PARP3 catalyse the transfer of a single ADP-ribose unit, mono (ADP-ribose) (MAR). In contrast, poly-ADP-ribosyltransferases, including human PARP1, PARP2, Tankyrases, plant PARP2, and bacterial PARPs, synthesise ADP-ribose polymers PAR [^36,55,63,64^]. Although, subtle sequence and structural differences within catalytic domains contribute to these distinct outputs [^52,62,63,65,66^]. It remains unclear whether regulatory domains also influence catalytic specificity, allosteric activation, and the determination of mono-versus poly-ADP-ribosylation outcomes.

Here, we hypothesised that regulatory domains, either in whole or in part, govern catalytic specificity, DNA-dependent signalling, allosteric activation, HD-domain reorganisation, and the ultimate enzymatic outcome of mono- or poly-ADP-ribosylation through conserved communication networks with catalytic domains. To test this hypothesis, we established a biochemical reconstitution platform using recombinant regulatory and catalytic domains derived from evolutionarily distant mammalian, plant, and bacterial ARTs (Fig. 1C). Regulatory–CAT domain complementation experiments were performed between the mono-ADP-ribosyltransferases (MART) human PARP3 and the poly-ADP-ribosyltransferases (PART) human PARP1 (hPARP1), human PARP2 (hPARP2), Arabidopsis PARP2 (atPARP2), and the bacterial *Herpetosiphon aurantiacus* PARP (HAPARP) (Fig. 1C). These non-natural assemblies enabled direct examination of the general and specific combining abilities of regulatory and catalytic domains and allowed us to determine how regulatory modules influence catalytic activation, allosteric communication, and catalytic plasticity in response to phosphorylated and unphosphorylated single-strand (SSB) and double-strand (DSB) DNA breaks. Using SDS-PAGE-based PARylation and NAD^+^ hydrolysis assays, measurements of auto- and cross-ADP-ribosylation, and quantitative biolayer interferometry (BLI) analyses of regulatory–CAT domain interactions, we demonstrate that conserved allosteric communication pathways can be functionally transferred between evolutionarily divergent ART family members. Our findings reveal unexpected plasticity in regulatory–catalytic coupling and provide mechanistic insights into how modular protein architectures evolve new biological functions while preserving ancestral catalytic frameworks.

Here, we note that in the literature, ADP-ribosyltransferases (ARTs) are classified into mono-ADP-ribosyltransferases (MARTs) and poly-ADP-ribosyltransferases (PARTs), based on whether they transfer a single ADP-ribosyl unit or multiple ADP-ribosyl units, respectively; both classes are collectively referred to as Poly (ADP-ribose) polymerases (PARPs). In this study, we employed HD-domain-containing, DNA-break-dependent, allosterically activated MART (hPARP3) and PARTs (hPARP1, hPARP2, atPARP2, and HaPARP) in an in *vitro* reconstituted chimeric assembly to investigate (i) the cross-complementation/coupling ability between the regulatory and catalytic domains of MART and PARTs, and (ii) the allosteric activation of the catalytic (CAT) domain via recognition of allosteric ligands — such as phosphorylated double-strand break DNA (pDSB-DNA) and phosphorylated single-strand break DNA (pSSB-DNA) — by the regulatory domains.

## Results

### A MARylating regulatory domain cross-complements PARylating catalytic domains and restores the stimulation of DNA-dependent PARylation activity

MARTs and PARTs have highly conserved CAT domains, whereas their regulatory domains have evolved extensively (Fig. 1A) [^67^]. It remains unknown whether regulatory domains from MARTs can functionally substitute for those of PARTs. We therefore tested whether the regulatory domain of human PARP3 (REG_hPARP3_), a MARylating/MART enzyme, could cross-complement the catalytic domains of the PARylating/PART enzymes human PARP1 (CAT_hPARP1_), human PARP2 (CAT_hPARP2_), and Arabidopsis PARP2 (CAT_atPARP2_) (Fig. 1C). Given that phosphorylated DNA double-strand breaks (pDSB-DNA) have been shown to activate PARP2-family and PARP3 enzymes, these DNA substrates were included to assess DNA-dependent allosteric activation and signal transmission between regulatory and catalytic modules.

To examine functional complementation, in vitro PARylation assays were performed using equimolar amounts of REG_hPARP3_ and the catalytic domain of PARylating PARP in the presence or absence of pDSB-DNA (Supplementary Table 1) (Fig. 1D–F). SDS-PAGE-based PARylation assays revealed that REG_hPARP3_ alone could complement both CAT_hPARP2_ and CAT_atPARP2_, resulting in trans-PARylation of REG_hPARP3_ even in the absence of DNA (lane 7, Fig. 1D,E). In contrast, no detectable complementation was observed with CAT_hPARP1_ under DNA-free conditions (lane 7, Fig. 1F). Remarkably, the addition of pDSB-DNA enabled REG_hPARP3_ to complement all three catalytic domains, resulting in robust cis- and trans-PARylation of both the regulatory and catalytic components (lane 8, Fig. 1D-F). Furthermore, PARylation levels were substantially higher in the presence of DNA breaks than in DNA-free reactions, indicating efficient DNA-dependent activation of the chimeric regulatory–catalytic assemblies.

To quantitatively validate these observations, NAD^+^ consumption was measured using a chemical assay that determines residual NAD^+^ levels following catalysis (Fig. 2A-C). Consistent with the SDS-PAGE analyses, isolated catalytic domains exhibited only weak intrinsic activity, consuming less than 50 pmol NAD^+^, indicating that stand-alone CAT domains are largely inactive in the absence of regulatory input. The addition of REG_hPARP3_ increased basal PARylation activity approximately two-fold for all three catalytic domains, despite originating from a MARylating PARP family member. In the presence of DNA breaks, REG_hPARP3_ stimulated catalytic activity even further, resulting in approximately ∼10-fold, ∼15-fold, and ∼17-fold increases in NAD^+^ consumption by CAT_hPARP1_, CAT_hPARP2_, and CAT_atPARP2_, respectively.

**Fig. 2.**
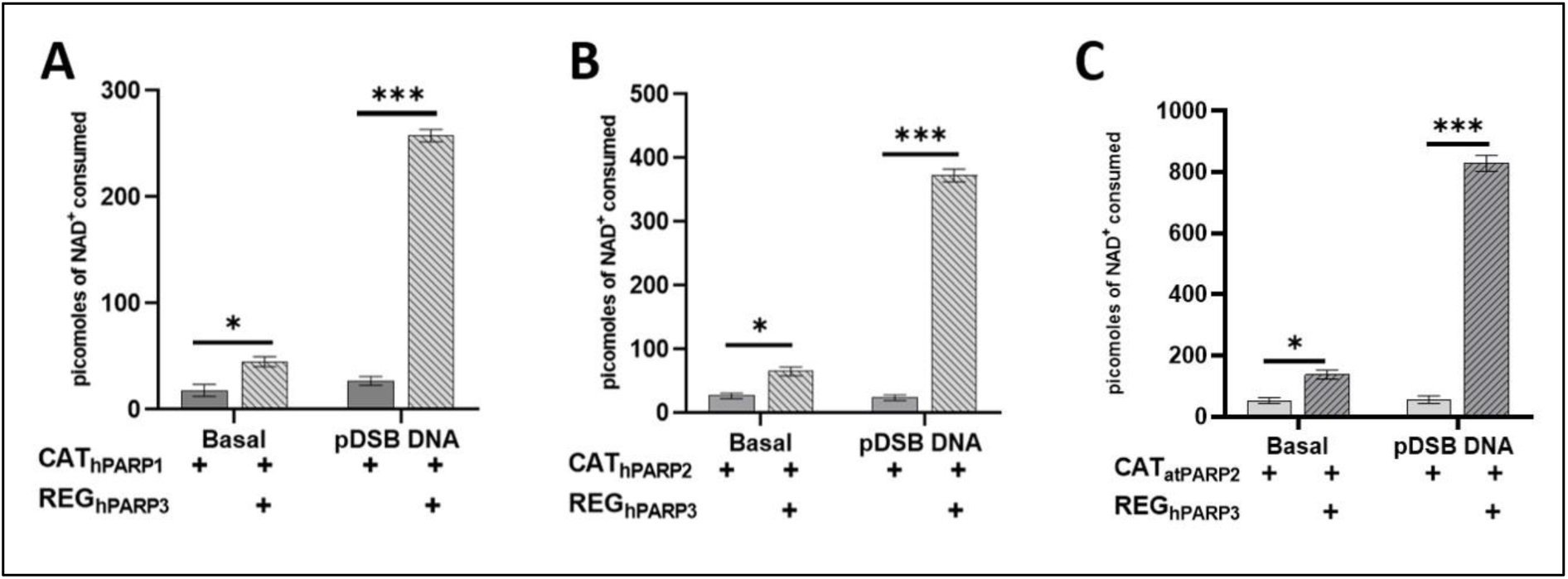
Chemical quantification of cross-complementation of the REG domain of MART and the CAT domain of PARTs to measure PARylation activity: NAD+ consumption by in vitro assembled. (A) CAT_hPARP1_, (B) CAT_hPARP2_, (C) CAT_atPARP2_, with REG_hPARP3_ in the presence of pDSB-DNA. Error bars show SEM, n = 3. The p-values were calculated using a 2-way ANOVA (where ns, *, **, *** indicate p > 0.05, < 0.05, <0.01, < 0.001, respectively).

These findings demonstrate that the regulatory domain of the MART enzyme PARP3 can functionally complement catalytic domains from evolutionarily distinct PARylating (PART) PARPs. The ability of REG_hPARP3_ to enhance both basal and DNA-dependent catalytic activity suggests that regulatory domains provide conserved allosteric functions that are not restricted by mono- or poly-ADP-ribosylation classification. The pronounced stimulation observed upon DNA binding further supports the existence of a conserved signal-relay mechanism through which regulatory modules communicate with catalytic domains to promote PARylation. Collectively, these results indicate that regulatory and catalytic domains of PARP-family enzymes retain a remarkable degree of functional interoperability despite substantial evolutionary divergence.

### MART-REG_hPARP3_ complements PARylating catalytic domains in response to phosphorylated single-strand DNA breaks

Previous studies have demonstrated that human PARP2 participates in the single-strand break (SSB) repair pathway and that both hPARP2, a PARylating PARP, and hPARP3, a MARylating PARP, are activated by phosphorylated single-strand break DNA (pSSB-DNA) substrates [^51^]. Given the ability of REG_hPARP3_ to complement PARylating catalytic domains in response to double-strand DNA breaks, we next investigated whether the MART regulatory domain of hPARP3 could similarly activate the catalytic domains of hPARP1, hPARP2 and atPARP2 in the presence of pSSB-DNA (Supplementary Table 1). This experiment was designed to determine whether the functional compatibility between regulatory and catalytic modules extends across distinct DNA damage signals and whether pSSB-DNA can promote allosteric communication between MART-derived regulatory domains and PARylating catalytic domains.

To assess regulatory–catalytic coupling for catalytic activity, in vitro PARylation assays were performed using equimolar amounts of REG_hPARP3_ and either CAT_hPARP1_, CAT_hPARP2_ or CAT_atPARP2_ in the presence or absence of pSSB-DNA (Supplementary Table 1) (Fig. 3A–C). Consistent with our observations using the pDSB-DNA allosteric activator, the REG_hPARP3_ complemented both CAT_hPARP2_ and CAT_atPARP2_ in the absence of pSSB-DNA, resulting in cross-PARylation of REG_hPARP3_ (lane 7, Fig. 3B,C). In contrast, no detectable PARylation was observed with CAT_hPARP1_ under DNA-free conditions (lane 7, Fig. 3A). Upon addition of pSSB-DNA, REG_hPARP3_ efficiently complemented all three catalytic domains, resulting in both cis- and trans-PARylation of REG_hPARP3_ and the corresponding catalytic domains (lane 8, Fig. 3A–C). These findings demonstrate that pSSB-DNA is sufficient to promote productive communication between REG_hPARP3_ and PARylating catalytic domains from both mammalian and plant PARPs.

**Figure 3.**
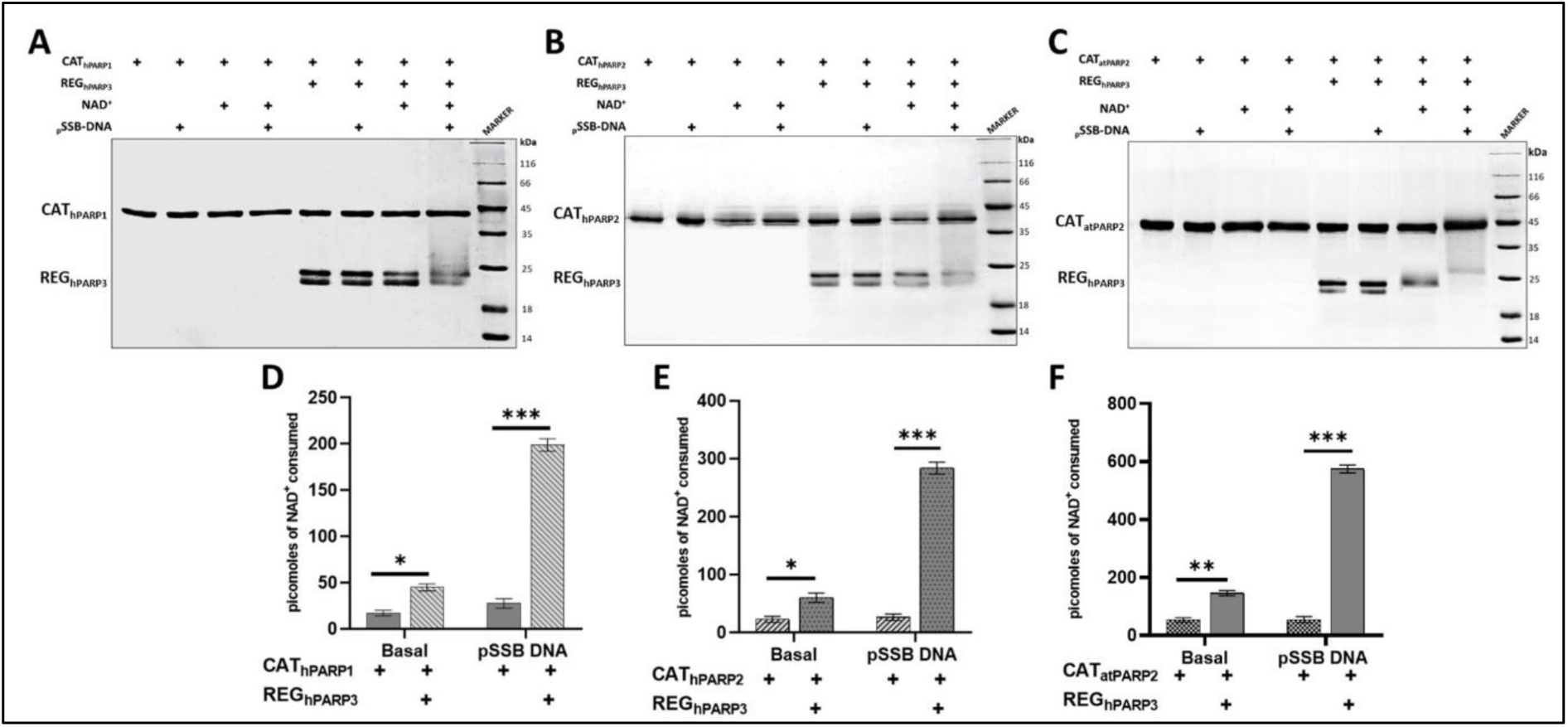
Qualitative and quantitative PARylation assays of cross-complementation of the REG domain of MART and the CAT domain of PARTs to measure the DNA break-dependent catalytic activity of chimeric assemblies. SDS-PAGE-based PARylation assay for cross-complementation of (A) CAT_hPARP1_ (B) CAT_hPARP2_, (C) CAT_atPARP2_, in the presence of pSSB-DNA, by REG_hPARP3_. NAD^+^ consumption measurement to quantify the PARylation activity of in vitro assemblies of (D) CAT_hPARP1_, (E) CAT_hPARP2_, (F) CATatPARP2, and REGhPARP3 in the presence of pSSB-DNA. Error bars show SEM, n = 3. The p-values were calculated using a 2-way ANOVA (where ns, *, **, *** indicate p > 0.05, < 0.05, <0.01, < 0.001, respectively).

To quantitatively evaluate catalytic activity, residual NAD^+^ was measured following the reactions using a chemical NAD^+^ consumption assay (Fig. 3D–F). The quantitative measurements corroborated the SDS-PAGE analyses and further revealed effects that were not readily detectable by gel-based methods. In the absence of DNA, REG_hPARP3_ enhanced the basal catalytic activity of all three catalytic domains, resulting in approximately 3-5-fold increases in DNA-independent activity for all-CAT_hPARP1_, CAT_hPARP2_ and CAT_atPARP2_, compared to CAT domains alone (Fig. 3D-F). In the presence of pSSB-DNA, REG_hPARP3_ further stimulated catalytic activity, resulting in approximately ∼10-fold, ∼12-fold and ∼15-fold increases in DNA-dependent NAD^+^ consumption for CAT_hPARP1_, CAT_hPARP2_ and CAT_atPARP2_, respectively (Fig. 3D-F). Together, these data demonstrate that REG_hPARP3_ can efficiently complement all three PARylating catalytic domains under pSSB-DNA activating conditions.

These results demonstrate that the MART-derived regulatory domain of hPARP3 can functionally engage catalytic domains of evolutionarily distinct PARylating enzymes in response to single-strand DNA break signals. REGh_PARP3_ strongly complements CAT_hPARP2_ and CAT_atPARP2_ even under DNA-independent conditions, whereas complementation of CAT_hPARP1_ is comparatively weak in the absence of DNA. Importantly, pSSB-DNA markedly enhances complementation of all catalytic domains, indicating that DNA-induced allosteric activation can be transmitted across non-cognate regulatory–catalytic interfaces. However, compared with the pDSB-DNA-mediated stimulation, the pSSB-DNA-mediated stimulation is less potent in activating the reconstituted complexes, suggesting that the efficiency of signal transmission and catalytic activation might depend on the nature of the DNA damage allosteric activator. Collectively, these findings support the existence of a conserved, interchangeable allosteric communication mechanism within the PARP family, while also highlighting ligand-specific allosteric differences in activation efficiency.

### Physical coupling of a MARylating regulatory domain and a PARylating catalytic domain from different species underlies the DNA-breaks-dependent cross-coupling for the catalytic activity

The finding that REGh_PARP3_, the regulatory domain of the mono-ADP-ribosyltransferase hPARP3, most efficiently stimulated the catalytic activity of CATatPARP2, the catalytic domain of the poly-ADP-ribosyltransferase atPARP2, was unexpected given their distinct regulatory domains, enzymatic outputs, and evolutionary origins. While hPARP3 catalyses mono-ADP-ribosylation and atPARP2 synthesises poly(ADP-ribose), the strong catalytic activation observed upon their complementation suggests that these domains may retain conserved interaction interfaces capable of supporting productive allosteric communication. We therefore sought to determine whether direct physical interactions between REGh_PARP3_ and CAT_atPARP2_ underlie their functional compatibility.

To test this hypothesis, Bio-Layer Interferometry (BLI) was employed using biotinylated REG_hPARP3_ immobilised on streptavidin biosensors and purified CAT_atPARP2_ as the analyte. BLI analyses revealed a direct interaction between the two domains with an equilibrium dissociation constant (K_D_) of approximately 14 μM (Fig. 4A). These results demonstrate that a regulatory domain derived from a MARylating PARP can directly associate with a catalytic domain derived from a PARylating PARP, providing a molecular basis for the efficient catalytic complementation observed in the PARylation and NAD^+^ consumption assays.

**Figure 4.**
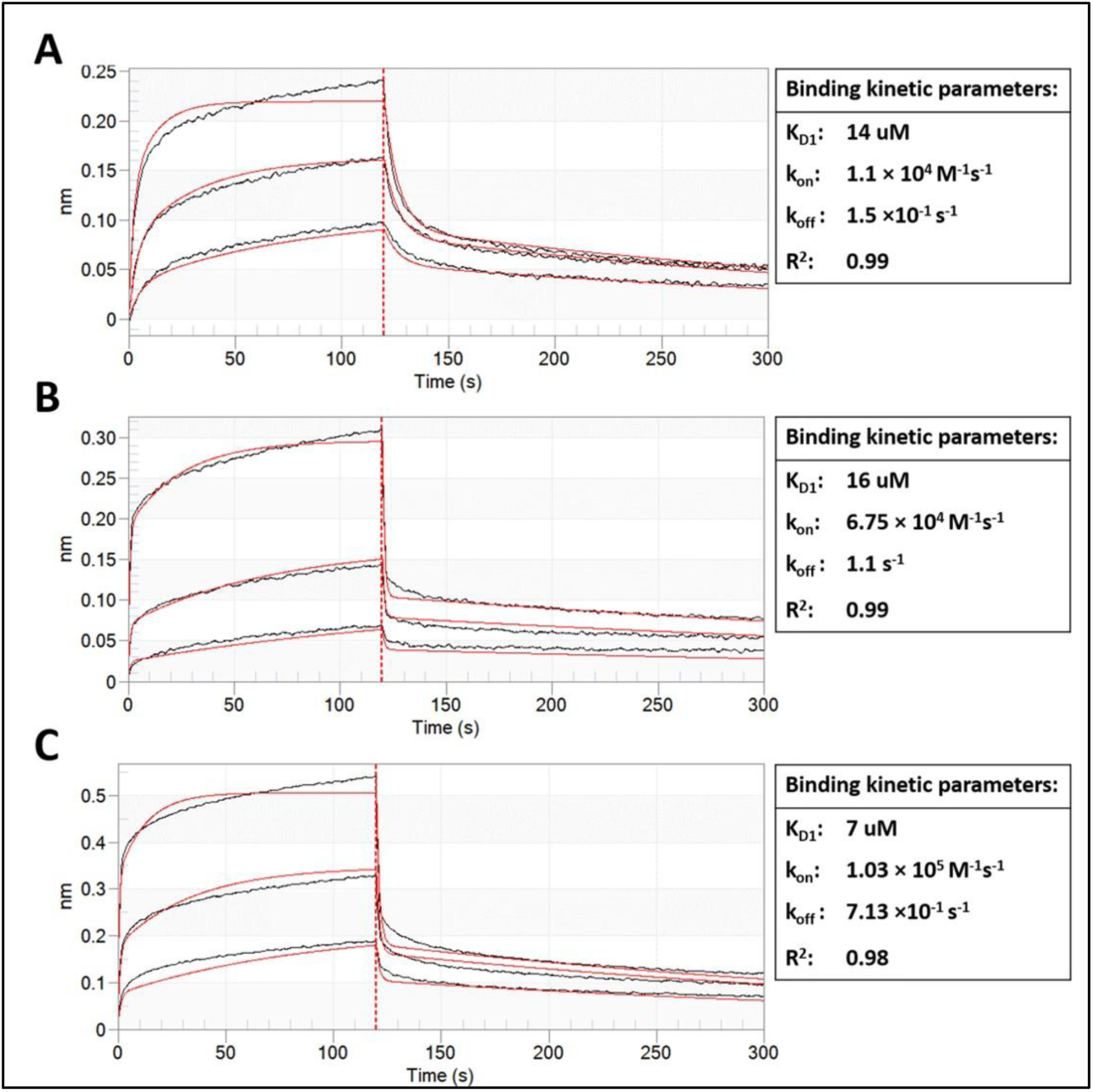
Bio-Layer Interferometry (BLI) binding studies between the REG domain of MART and the CAT domain of PART. (A) BLI sensogram of CAT_atPARP2_ for binding to REG_hPARP3_. Where REG_hPARP3_ was biotinylated and immobilised on the biosensor. The data were fit globally to the 1:1 binding model; The processed experimental data curves are in black; model fit curves from the 1:1 global analysis are shown in red. (B) BLI sensogram of REG_hPARP3_ binding to pDSB-DNA. (C) BLI sensogram of a 1:1 molar ratio of REG_hPARP3_-CAT_atPARP2_ complex binding to pDSB-DNA. The data were fit globally to the 2:1 binding model for B and C. The processed experimental data curves are in black; model fit curves from the 2:1 global analysis are shown in red. The binding kinetics data are given in the inset on the right of each sensogram. The R^2^ indicates the statistical confidence of the global fit of the experimental data to the model.

Because DNA binding is the initiating event in the allosteric activation of DNA-dependent PARPs, we next examined the interaction between REG_hPARP3_ and phosphorylated double-strand break DNA. BLI measurements showed that REG_hPARP3_ binds DSB-DNA with a binding affinity of approximately 16 μM (Fig. 4B), confirming its intrinsic DNA-recognition capability. We then investigated whether association with CAT_atPARP2_ affects the DNA-binding activity of REG_hPARP3_. Strikingly, the REG_hPARP3_–CAT_atPARP2_ complex exhibited an approximately 2-fold higher affinity for DNA than REG_hPARP3_ alone (Fig. 4B, C). Thus, formation of the heterologous regulatory–catalytic complex enhances DNA recognition, indicating functional cooperativity between the two domains.

These findings provide direct biophysical evidence that efficient complementation between REG_hPARP3_ and CAT_atPARP2_ is mediated by physical inter-domain interactions and cooperative DNA binding. The enhanced affinity of the REG_hPARP3_–CAT_atPARP2_ complex for DNA suggests that catalytic domain association stabilises a DNA-bound state that is more competent for allosteric activation. Importantly, these observations demonstrate that a regulatory module originating from a mono-ADP-ribosyltransferases can productively communicate with a catalytic domain that generates poly(ADP-ribose), indicating that the molecular framework governing regulatory–catalytic signal transmission is conserved despite divergence in enzymatic output (Fig. 1B). The ability of a MARylating regulatory domain to enhance activation of a PARylating catalytic domain further supports the existence of a common allosteric communication network shared across the ART superfamily (Fig. 1A,B). Because human PARP3 and Arabidopsis PARP2 are evolutionarily distant proteins, this observation raised the intriguing possibility that conserved interaction surfaces that enable regulatory–catalytic communication has been maintained across species boundaries. Future structural and biophysical studies will be required to define the molecular interfaces and conformational transitions that enable this unexpected cross-class and cross-species regulatory compatibility.

### PARylating PARP regulatory domains of paralog, hPARP2, fail to functionally complement the catalytic domain of MARylating PARP, hPARP3, despite conserved regulatory and catalytic domains

Our previous experiments revealed an unexpected degree of functional interoperability between the regulatory domain of the mono-ADP-ribosyltransferase hPARP3 (REG_hPARP3_) and catalytic domains derived from evolutionarily distant poly-ADP-ribosyltransferases. REG_hPARP3_ efficiently complemented the catalytic domains of human and plant PARPs, stimulated DNA-dependent catalytic activity, and directly interacted with CAT_atPARP2_; DNA further enhanced complex formation. These findings suggested that the fundamental allosteric pathway linking DNA-break recognition, regulatory-domain signalling, HD-domain remodelling, and catalytic activation remains conserved across divergent PARP-family enzymes. However, whether this functional compatibility is bidirectional remains unknown. Specifically, it was unclear whether regulatory domains from PARylating PARPs could similarly activate the catalytic domain of the MARylating enzyme hPARP3. Addressing this question is important for understanding whether regulatory–catalytic interoperability is a general property of ADP-ribosyltransferases or is constrained by the catalytic specialization of individual family members.

To investigate this possibility, we established a complementation assay that directly monitors MARylation. Recent advances in MARylation detection, including highly sensitive MAR-specific antibodies, have enabled reliable detection of mono-ADP-ribosylated proteins by immunoblotting. We therefore adapted our in vitro complementation assay by coupling it with western blot (WB) analysis using HRP-conjugated anti-MAR antibodies [^68^]. Following MARylation reactions, samples were resolved by SDS-PAGE, transferred to PVDF (polyvinylidene fluoride) membrane, and probed for MARylated proteins. Full-length hPARP3 was included as a positive MARylation control, while identical Coomassie-stained SDS-PAGE gels served as loading controls for each western blot (Fig. 5A, lane 10, Supplementary Fig. 2A).

**Figure 5.**
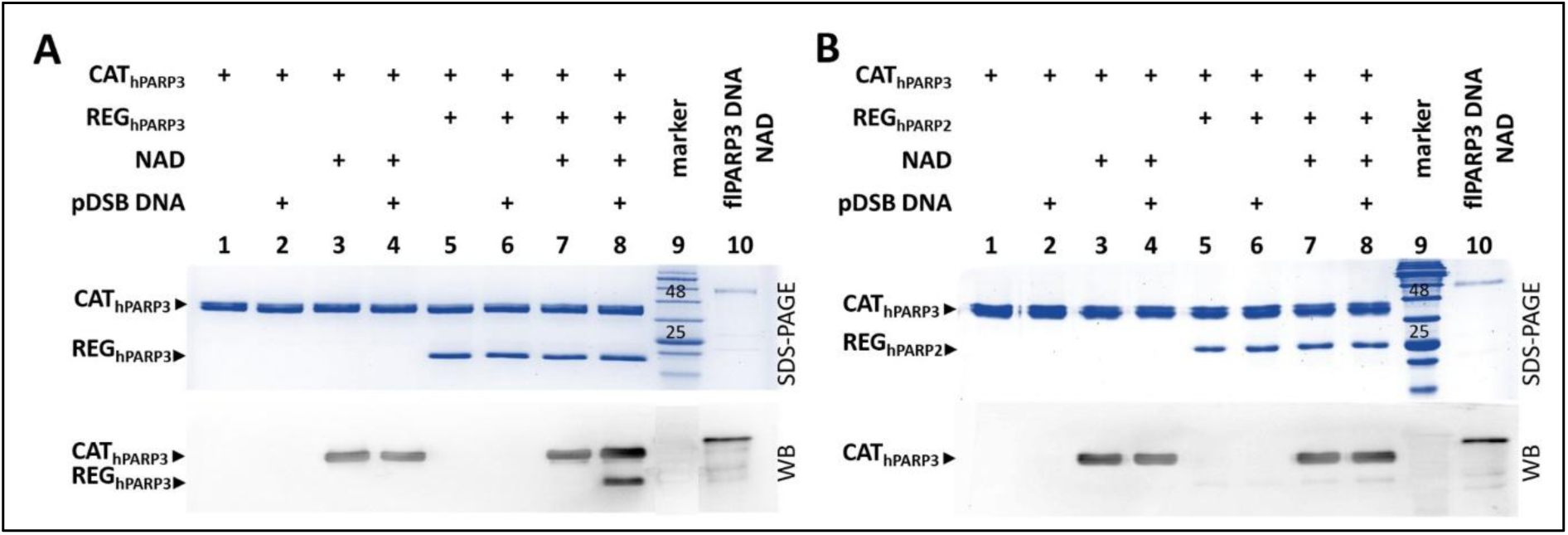
Western blot analysis of MARylation activity of biochemically reconstituted CAT domain of hPARP3 by the REG domain of hPARP2. (A) Cis-complementation of CAT_hPARP3_ by REG_hPARP3_ for pDSB-dependent MARylation activity. (B) Cross-complementation of CAT_hPARP3_ by REG_hPARP2_ for pDSB-dependent MARylation activity. flPARP3 is used as a MARylation-positive control in all gels (lane 10). HRP-tagged anti-MAR antibody is used on all blots to detect MARylation. The upper blue gels in both panels are control SDS-PAGE gels. Each western blot experiment was repeated three times.

We first examined the cognate REG_hPARP3_–CAT_hPARP3_ pair. Western blot analyses demonstrated that isolated CAT_hPARP3_ exhibits weak basal MARylation activity, regardless of DNA damage (Fig. 5A, lanes 3 and 4, Supplementary Fig. 2A). In contrast, efficient MARylation of both REG_hPARP3_ and CAT_hPARP3_ was observed only in the presence of DNA breaks (Fig. 5A, compare lanes 7 and 8, Supplementary Fig. 2A), indicating that productive complementation requires DNA-dependent activation. The presence of REG_hPARP3_ alone was insufficient to stimulate detectable MARylation in the absence of DNA (Fig. 5A, lanes 3 and 4), notably, unlike the PARylating systems examined above and previously reported by our group [^37^]. We also tested whether regulatory domains from PARylating enzymes could complement the catalytic domain of hPARP3. REG_hPARP2_ was combined with CAT_hPARP3_ and analysed under identical conditions. Identical to the above results, isolated CAT_hPARP3_ exhibited basal MARylation activity regardless of the DNA (Fig. 5B, lanes 3 & 4, Supplementary Fig. 2B). Western blot analyses failed to detect DNA-dependent MARylation of the regulatory domain of hPARP2 by the CAT_hPARP3_ (Fig. 5B, lane 8, Supplementary Fig. 2B) in the presence of REG_hPARP2_. Consistent with these observations, Coomassie-stained SDS-PAGE loading controls showed no evidence of PARylation-associated band smearing or loss of protein signal, confirming that the observed lack of MARylation was not due to PARylation, protein instability or differential loading. The faint MARylation bands observed for REG_hPARP2_ (Fig. 5B, lanes 7, 8) were comparable to those detected in the absence of NAD⁺ and DNA (Fig. 5B, lanes 5, 6), suggesting that they represent nonspecific background signals rather than bona fide MARylation. Therefore, these bands were considered negligible. Further extensive quantitative analysis would be required to determine whether REG_hPARP2_ is MARylated by CAT_hPARP3_.

### Regulatory domains of PARTs fail to functionally complement the catalytic domain of the MART despite conserved regulatory–catalytic communication in PARPs

We also tested whether the regulatory domains of other PARP3 orthologs, REG_atPARP2_ and REG_HAPARP_, could complement CAT_hPARP3_ for DNA-break-dependent MARylation. We have chosen the regulatory domains of the bacterial Herpetosiphon aurantiacus PARP (HAPARP) and atPARP2 for the following reasons: Both hPARP3 and HAPARP have identical domain architecture (Fig. 1A), the catalytic domain of atPARP2 exhibited robust, maximal catalytic activity when coupled with the regulatory domains of hPARP3 with the allosteric activators pDSB and pSSB (Fig. 2 and Fig. 3). Consistent with the above results, isolated CAT_hPARP3_ exhibited basal MARylation activity regardless of the DNA (Fig. 6, lanes 3 & 4, Supplementary Fig. 2C,D). However, as with the above results for REG_hPARP2_, both REG_atPARP2_ and REG_HAPARP_ fail to complement CAT_hPARP3_ despite the presence of DNA (Fig. 6A and B, lane 8; Supplementary Fig. 2C,D).

**Figure 6.**
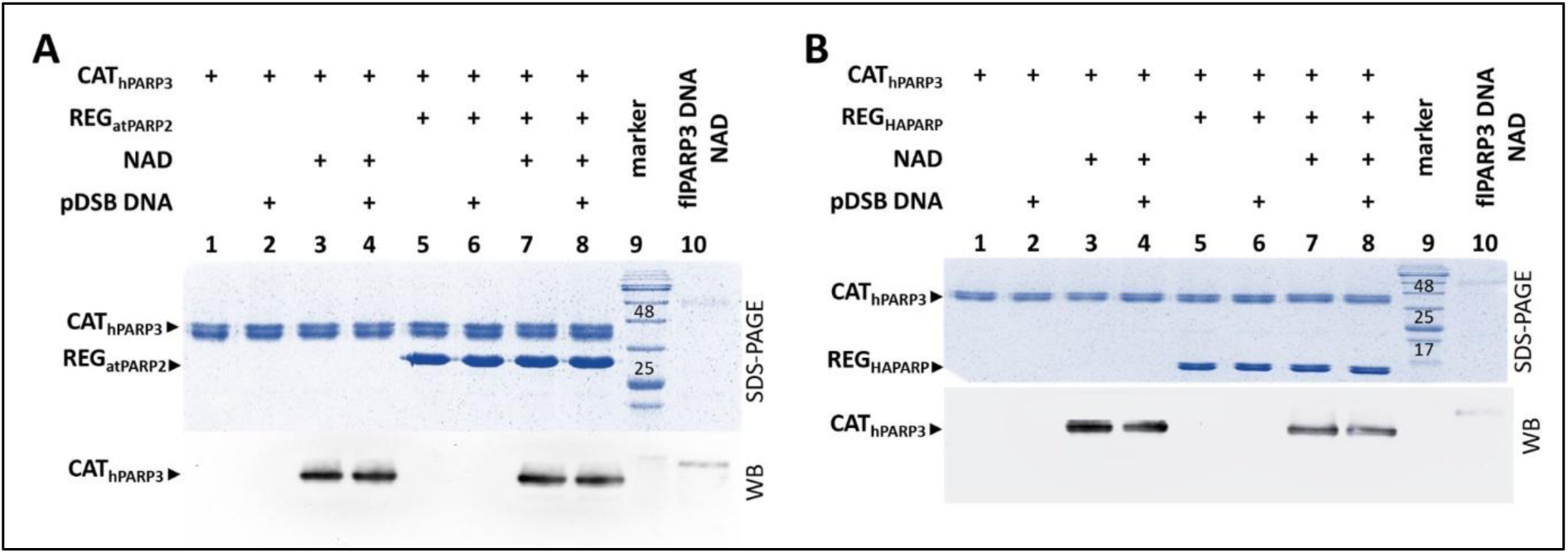
Western blot analysis of MARylation activity of biochemically reconstituted chimeric assemblies of the CAT domain of MART and the REG domains of PARTs. Cross-complementation of regulatory domains of (A) atPARP2 (REG_atPARP2_) and (B) HAPARP (REG_HAPARP_) for pDSB-dependent MARylation activity. The flPARP3 is used as a MARylation positive control in all gels (lane 10). HRP-tagged anti-MAR antibody is used on all blots to detect MARylation. The upper blue gels in both panels are unprocessed control SDS-PAGE gels. Each western blot experiment was repeated three times.

These results demonstrate a striking asymmetry in regulatory–catalytic compatibility within the PARP family. Whereas the regulatory domain of the MARylating enzyme hPARP3 can efficiently activate catalytic domains from multiple PARylating enzymes, regulatory domains from PARylating PARPs are unable to activate the catalytic domain of hPARP3. This finding indicates that the conserved allosteric communication machinery identified in PARylating PARPs is not universally interchangeable. Instead, activation of the hPARP3 catalytic domain appears to require specific regulatory-domain features absent in PARylating PARPs. The strict dependence on DNA breaks observed for the cognate REG_hPARP3_–CAT_hPARP3_ pair (Fig. 5A, Lanes 7 & 8) further suggests that hPARP3 has evolved a more tightly controlled activation mechanism, consistent with its specialised role in DNA damage signalling. Collectively, these findings support a model in which regulatory domains can broadly communicate with PARylating catalytic domains, but activation of the MARylating hPARP3 catalytic domain requires highly specific regulatory inputs that have been selectively maintained during evolution.

## Discussion

### General and specific combining ability reveal distinct modes of regulatory–catalytic communication in PARPs

The present study demonstrates that regulatory and catalytic domains from evolutionarily divergent ADP-ribosyltransferases retain an unexpected capacity for functional interoperability. Extending our previous findings on cross-complementation among DNA-damage-responsive PARylating PARPs [^37^], we show that the regulatory domain of the mono-ADP-ribosyltransferases hPARP3 can functionally activate catalytic domains from hPARP1, hPARP2, and atPARP2 in response to both phosphorylated double-strand and single-strand DNA breaks. These observations indicate that the molecular machinery required for DNA-break recognition, allosteric signal propagation, HD-domain remodelling, and catalytic activation remains sufficiently conserved (Fig. 1B) to permit productive communication between domains that have evolved in different proteins and even different kingdoms of life.

Our findings further suggest that regulatory–catalytic compatibility can be interpreted through the concepts of general combining ability (GCA) and specific combining ability (SCA). REG_hPARP3_ exhibits characteristics of a broadly compatible regulatory module capable of activating multiple heterologous PARylating catalytic domains, analogous to high GCA observed in breeding and combinatorial engineering systems [^69–73^]. In contrast, the catalytic domain of hPARP3 exhibits strong SCA, activating only in the presence of its cognate regulatory domain and failing to complement with regulatory domains derived from PARPylating enzymes. Thus, although PARylating catalytic domains appear permissive to regulatory-domain substitution, the MARylating CAT_hPARP3_ imposes substantially more stringent requirements for productive allosteric communication.

This asymmetry has important mechanistic implications. The ability of REG_hPARP3_ to activate multiple PARylating catalytic domains suggests that the regulatory domain contains conserved signalling features that can engage diverse catalytic partners. Conversely, the inability of REG_hPARP1_, REG_hPARP2_, REG_atPARP2_, or REG_haPARP_ to activate CAT_hPARP3_ implies that activation of the MARylating catalytic domain depends on highly specific molecular interactions that have been maintained during evolution. These observations suggest that catalytic specialisation may be accompanied by increased stringency in regulatory-domain recognition and allosteric coupling.

### PARPs as a model system for investigating the evolution of modular protein communication

Modular proteins are pervasive throughout biology, including kinases [^74^] (Supplementary Fig 1), GPCRs [^22–24,75^], ubiquitin ligases [^76–79^], transcription factors [^80,81^], and transmembrane receptors [^18,19^]. However, PARPs may represent one of the most powerful systems for investigating how regulatory and catalytic domains co-evolve, as they combine a highly conserved catalytic fold with extraordinary regulatory diversity [^35,67^]. Hundreds of ART-family proteins, including Mono-ADP-ribosyltransferases and Poly-ADP-ribosyltransferases, have been identified across mammals, plants, fungi, insects, bacteria, archaea, and viruses [^82,83^], where they participate in remarkably diverse biological processes ranging from DNA repair and genome maintenance to immunity, mitochondrial homeostasis, development, host–pathogen interactions, and stress responses [^47–49,84^].

Previous studies, including our ACS Chemical Biology report on PARP regulatory–catalytic cross-complementation and investigations of Arabidopsis PARP family members, demonstrated that catalytic activation can be restored through non-cognate domain combinations [^38^, ^85^]. The present work extends this concept beyond PARylating enzymes by incorporating a MARylating PARP and revealing fundamental differences in combining behaviour. Together, these studies support a model in which ART-family proteins contain a mixture of highly conserved and highly specialised allosteric interfaces. Successful complementation events likely reflect preservation of ancestral communication networks, whereas failed complementation events reveal evolutionary innovations that generate signalling specificity.

Importantly, these observations suggest that qualitative complementation (“yes/no”) and quantitative complementation (degree of activation) should be viewed as distinct biological properties. A comprehensive understanding of PARP evolution will therefore require systematic measurement of both parameters across large numbers of regulatory-catalytic domain combinations. Such datasets could reveal whether particular domains exhibit broad GCA, strict SCA, or intermediate compatibility profiles, and identify the structural features responsible for these behaviours.

### Toward a cross-complementation atlas of the ADP-Ribosyltransferase superfamily

One of the most exciting implications of this work is the possibility of creating a comprehensive cross-complementation database encompassing ARTs from multiple kingdoms of life. Such a resource would catalogue regulatory–catalytic compatibility, inter-domain binding affinities, catalytic outputs, allosteric ligand requirements, and DNA-dependent activation profiles across hundreds of proteins. Beyond simple classification, this database would establish a functional map of ART-family communication networks.

Future studies should expand beyond DNA-break-dependent PARPs and include activation by structured RNA [^58,86^], G-quadruplex DNA [^60,87^] proteins [^88–91^] post-translational modifications [^92,93^] and poly (ADP-ribose] itself [^41^]. Because many PARPs respond to distinct classes of allosteric ligands, assessing complementation across diverse activation contexts may reveal why some domains exhibit broad GCA whereas others display stringent SCA. Equally important will be biochemical, biophysical, and structural investigations aimed at defining the molecular basis of successful and unsuccessful complementation. Cryo-EM, X-ray crystallography, HDX-MS, NMR spectroscopy, molecular dynamics simulations, and co-evolutionary analyses should help identify the interaction surfaces and conformational transitions that govern regulatory–catalytic compatibility.

Such efforts may also enable the development of reverse-allostery studies. Whereas most current models focus on signal transmission from regulatory domains to catalytic centres, ligand binding within catalytic domains—including NAD^+^, PARP inhibitors, or ADP-ribosylation reaction intermediates—may reciprocally influence regulatory-domain conformations and DNA-binding properties. Understanding these bidirectional communication pathways could substantially reshape current models of PARP regulation and open new therapeutic opportunities.

### Engineering synthetic PARP systems for biology, medicine, and drug discovery

The broader significance of cross-complementation extends beyond PARP biology. The ability to identify compatible regulatory and catalytic modules raises the possibility of engineering synthetic PARP systems with programmable inputs and outputs. Analogous to the success of CAR-T receptors [^94^], synthetic Notch receptors [^95^], designer GPCRs [^96,97^], and engineered kinase circuits [^98^], artificial PARPs could be constructed by combining regulatory domains from one organism with catalytic domains from another. Such chimeric enzymes could function as biosensors, signal amplifiers, molecular switches, or logic gates that respond to specific DNA structures, proteins, RNAs, metabolites, or environmental stimuli.

Engineered PARP systems would provide powerful tools for studying DNA repair [^35^], apoptosis [^47^], chromatin regulation [^99^], innate immunity [^49^], liquid–liquid phase separation [^100^], stem-cell biology [^101^], development and differentiation [^101,102^], and PAR-mediated signalling [^103^]. By selectively rewiring allosteric communication pathways, investigators could directly test how ADP-ribosylation contributes to specific biological outcomes. Such systems could also serve as orthogonal signalling modules in synthetic biology and systems biology applications.

The implications for drug discovery are equally substantial. Current PARP inhibitors predominantly target the conserved NAD^+^-binding site, limiting selectivity across family members. Cross-complementation studies provide a framework for identifying conserved and species-specific allosteric interfaces that may be therapeutically targetable. Artificially reconstituted PARP systems could be adapted for high-throughput screening of compounds that selectively modulate DNA recognition, inter-domain communication, catalytic activation, or PAR-chain synthesis. Furthermore, patient-derived variants could be evaluated within complementation platforms to identify defects in allosteric signalling and generate functional biomarkers for diagnosis and precision medicine.

### Conclusion/Summary

In conclusion, our findings establish that regulatory–catalytic communication within the PARP family is simultaneously conserved and specialised. Whereas PARylating catalytic domains exhibit considerable compatibility with heterologous regulatory modules, the MARylating catalytic domain of hPARP3 requires highly specific regulatory inputs. These observations introduce the concepts of GCA and SCA into PARP biology and provide a conceptual framework for understanding the evolution of modular signalling proteins. More broadly, they suggest that systematic cross-complementation studies across the ART superfamily can generate a predictive atlas of allosteric communication, enabling both mechanistic discovery and the rational engineering of synthetic PARP systems for applications in basic research, biotechnology, diagnostics, and therapeutics.

## Methods

### Cloning, expression, and purification of REG and CAT domains of different PARP<u>s</u>

The hPARP1 (UNIPROT ID: P09874) cDNA cloned in pET28-a(+) was purchased from GenScript, USA. The hPARP2 isoform 2 (UNIPROT ID: Q9UGN5-2) cDNA cloned in pET28-a(+) was generously gifted by Dr. John Pascal, Université de Montréal. The haPARP (UNIPROT ID: A9B244) cDNA cloned in pET28-a(+) was generously gifted by Dr. Ivan Ahel, Sir William Dunn School of Pathology. atPARP2 (UNIPROT ID: Q11207) cDNA cloned in pET28-c(+) was generously gifted by Dr. Amangeldy K. Bisenbayev, al-Farabi Kazakh National University. hPARP3 isoform 1 (Uniprot ID: Q9Y6F1-1) cloned in the pSRF-DUET1 vector was purchased from Biotech Desk. Domain boundaries for REG_hPARP1_ (aa: 1-638), and CAT_hPARP1_ (aa: 662-1014) are as given in [^104^]. REG_hPARP2_ (aa: 1-210), CAT_hPARP2_ (aa: 222-570), REG_hPARP3_ (aa: 1-155), CAT_hPARP3_ (aa: 181-533), REG_atPARP2_ (aa: 1-255), CAT_atPARP2_ (aa: 286-637), REG_haPARP_ (aa: 1-88) were PCR amplified and cloned into bacterial expression vector pRSFDuet-1, between BamH1, Xho1 restriction sites. All the protein domains were expressed in E. coli Rosetta2 (DE3) and purified by using the same protocol used for the purification of hPARP1 as described previously [^104^].

### Assembly of phosphorylated and un-phosphorylated DSB and SSB DNAs

HPLC-purified lyophilized oligonucleotides and complementary strands (listed in Supplementary Table 1) were purchased from the oligo synthesis facility of W.M. Keck Foundation, Yale School of Medicine, USA. The oligos were dissolved and annealed as reported previously [^41^].

### SDS-PAGE PARylation assay

To perform the PARylation assay for complementation, the reactions were set up as described previously [^38^]. Essentially, 5 μM of CAT_hPARP1_/ CAT_hPARP2_/ CAT_hPARP3_/ CAT_atPARP2_ was incubated with 5 μM of REG_hPARP2_/ REG_hPARP3_/ REG_haPARP_/ REG_hPARP1_/ REG_atPARP2_ for 15 minutes at a room temperature to allow for possible protein-protein interaction to occur. It was followed by incubation with 5 μM of different DNA-break mimics (pSSB DNA/ pDSB DNA) (Supplementary Table 1) for 15 min, with subsequent addition of 1mM NAD^+^ in a PARylation buffer (10mM HEPES, pH 8, 25mM KCl, 5mM CaCl_2_, 5mM MgCl_2_, 2mM β-ME) at room temperature. The reactions were incubated for 30 mins and quenched by adding SDS-loading dye. The samples were run on 12-15% SDS-PAGE based on the protein sizes and analyzed after staining with Coomassie Brilliant Blue.

### Quantitative PARylation assay: chemical method

To assess the catalytic activity of chimeric PARP assemblies and the extent of PARylation, a quantitative NAD⁺ consumption assay using an acetophenone-KOH-formic acid-based fluorometric method was performed. In this assay, the quantity of the generated fluorescent product correlates linearly with the unused NAD^+^. We compared NAD⁺ consumption across REG_hPARP3_ + CAT_hPARP1/hPARP2/atPARP2_ combinations, with or without DNA substrates (e.g., pDSB and pSSB). Consumed NAD^+^ was calculated as described by [^40^]. The error bars indicate ±SEM. Experiments were performed in triplicate (n =3), and measurements were fitted using GraphPad Prism version 8.0 for windows (GraphPad Software). The p-values were calculated using a 2-way ANOVA where ns = P >0.05, * = p <0.05, ** = p <0.01, *** = p < 0.001.

### BLI binding studies

To measure the binding kinetics, BLI studies were carried out on an OctetK2 instrument (Pall Fortebio). For the domain-domain interaction binding study, REG_hPARP3_ was biotinylated as described previously [^38^]. Streptavidin (SA) Dip and Read™ biosensors (Pall Fortebio) were used to immobilize biotinylated REG_hPARP3_. To measure association kinetics, the REG_hPARP3_ immobilized biosensors were dipped into wells containing varying concentrations of CAT_atPARP2_ in gel filtration buffer for 120s, followed by dipping into wells containing buffer for 180s to measure the dissociation kinetics. One of the biosensors immobilized with biotinylated REG_hPARP3_ was dipped into a buffer during the association and dissociation phases for reference. The measurement was carried out at 25°C. The data were fit to a 1:1 model and analyzed using ForteBio Data Analysis 12 (Pall Fortebio).

For binding studies with DSB DNA, HPLC-purified 16-mer Biotinylated Double-strand DNA (Supplementary Table 1), which mimics pDSB-DNA, was purchased from the oligo synthesis facility of W.M. Keck Foundation, Yale School of Medicine, USA, and dissolved as described above. For analyzing binding kinetics, the BLI experiment was performed in the same manner as mentioned above, with the immobilization of the biotinylated-DSB-DNA onto the biosensors, which were then dipped into varying concentrations of REG_hPARP3_ or REG_hPARP3_-CAT_atPARP2_ complex. The data were fit to a 2:1 model and analyzed using ForteBio Data Analysis 12 (Pall Fortebio).

### Western Blot

MARylation reaction samples were prepared as described above for the PARylation reactions. For each western blot experiment, two SDS–PAGE gels were run in parallel. One gel was stained with Coomassie Brilliant Blue to assess protein loading, whereas the proteins from the second gel were transferred onto a PVDF membrane using 1× transfer buffer (25 mM Tris base, 0.1 M glycine, pH 8.3) at 90 V for 2 h. The membrane was blocked with TBST (25 mM Tris-HCl, pH 7.5, 150 mM NaCl and 0.05% Tween-20) containing 5% BSA for 30 min. The membrane was then washed three times with TBST for 5 min per wash and incubated with HRP-conjugated anti-MAR antibody (1:2000) ^68^for 1 h at room temperature. Following antibody incubation, the membrane was washed three times with TBST, and chemiluminescent signal was detected using ECL Substrate (32209, Thermo Fisher Scientific) and an iBright CL1000 imaging system with a 1s exposure.

## Data availability

The datasets generated during and/or analysed during the current study are available from the corresponding author on reasonable request.

## Acknowledgements

We thank the Department of Biotechnology at Indian Institute of Technology Hyderabad for providing infrastructure and a platform to perform the current research. We also thank Dr. John Pascal, Université de Montréal, for providing hPARP2 cDNA clone, Dr. Ivan Ahel for providing haPARP cDNA and Dr. Amangeldy K. Bisenbayev, al-Farabi Kazakh National University, for the atPARP2 cDNA clone.

## Author Contributions

These authors contributed equally: Makwana Milee and Khare Shambhavi. M.M. and K.S.: Investigation, methodology, data acquisition, curation, formal analysis, writing-review and editing; S.P.G.: Methodology; R.G.: Methodology, data acquisition; B.A.: Supervision, validation. E.R.: Conceptualisation, funding acquisition, supervision, formal analysis, writing-review and editing.

## Supplementary information

The online version contains supplementary material.

