## Supporting Figures and Tables for "Evolutionary plasticity of regulatory–catalytic coupling in mono- and poly-ADP-ribosyl transferases": Supplementary_24_09_26.docx

**Supplementary information:**

The online version contains supplementary material.

Correspondence and requests for materials should be addressed to Eerappa Rajakumara.

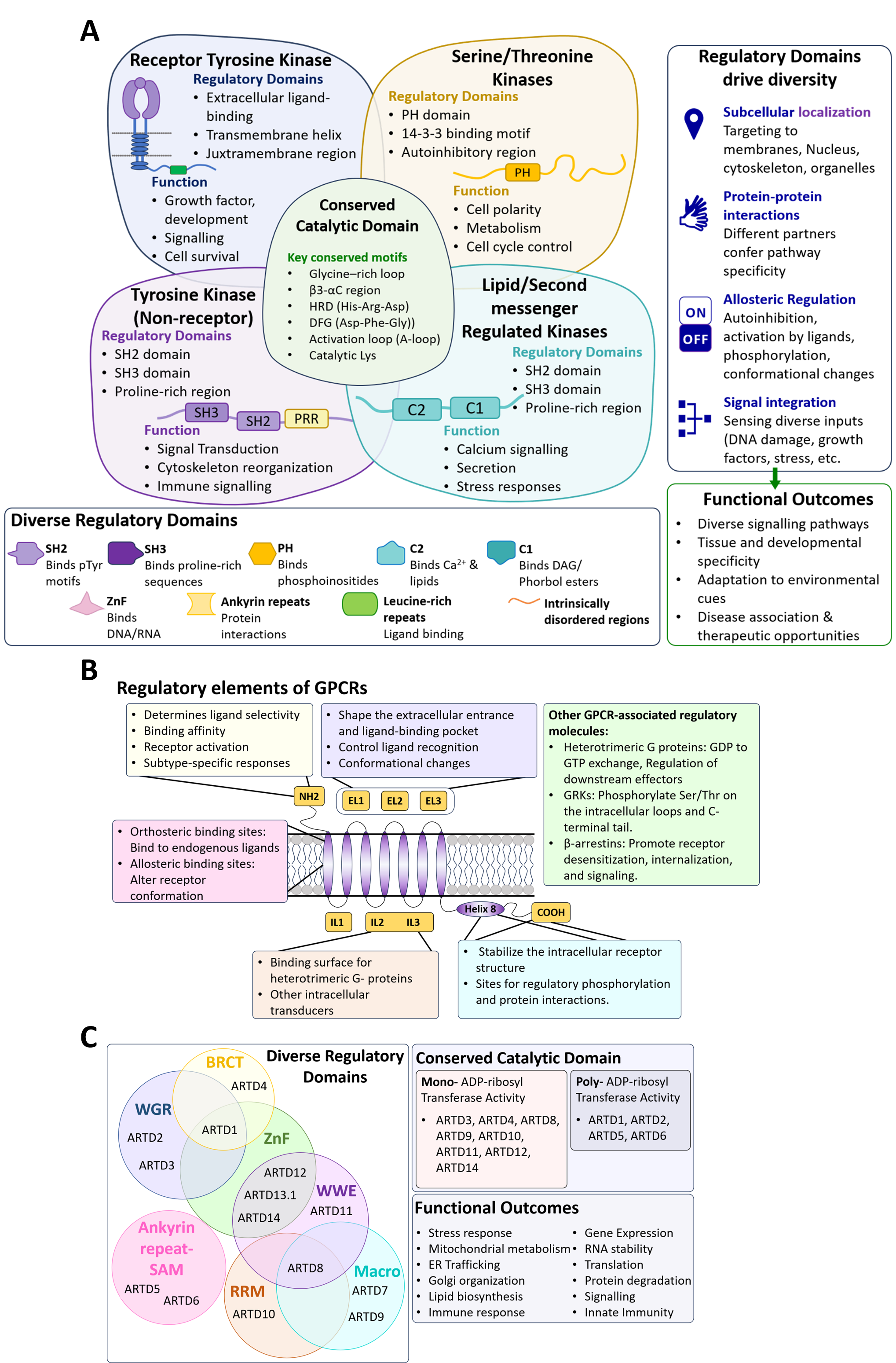

**Supplementary Figure 1. Modular Protein Architectures Enable Functional Diversification in** (A) protein kinases (B) GPCRs and (C) ADP-ribosyltransferases (ARTDs/ or PARPs).

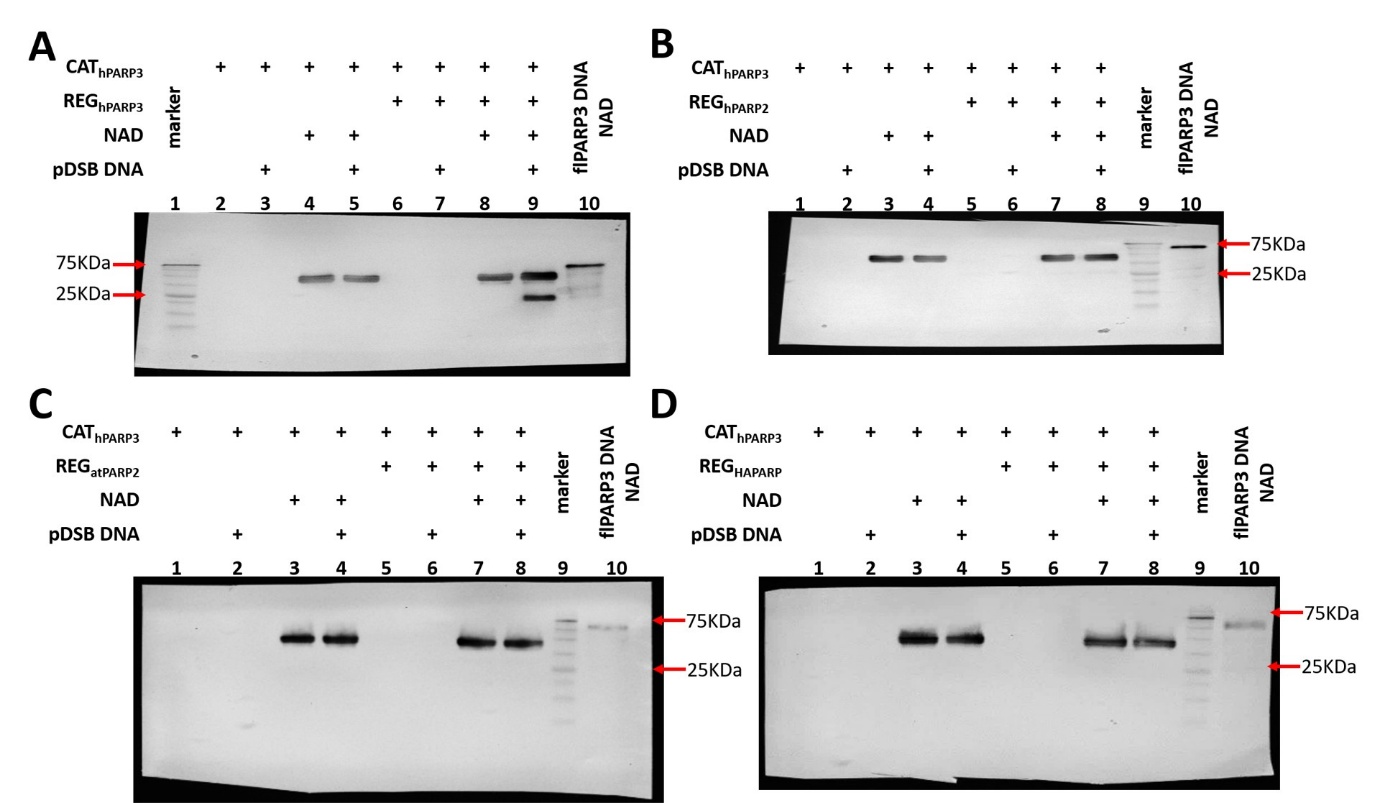

**Supplementary Figure 2. Unedited images of western blots using HRP-tagged anti-MAR antibodies.** Unprocessed membrane overlay scans corresponding to the cropped western blot images shown in Fig. 5 and 6, with full-length membranes including molecular weight markers (75 kDa, 25 kDa). Each membrane shows complementation of MART CAT_hPARP3_ by the indicated Regulatory domains (A) REG_hPARP3_, (B) REG_hPARP2_, (C) REG_atPARP2_ and (D) REG_HAPARP,_ respectively.

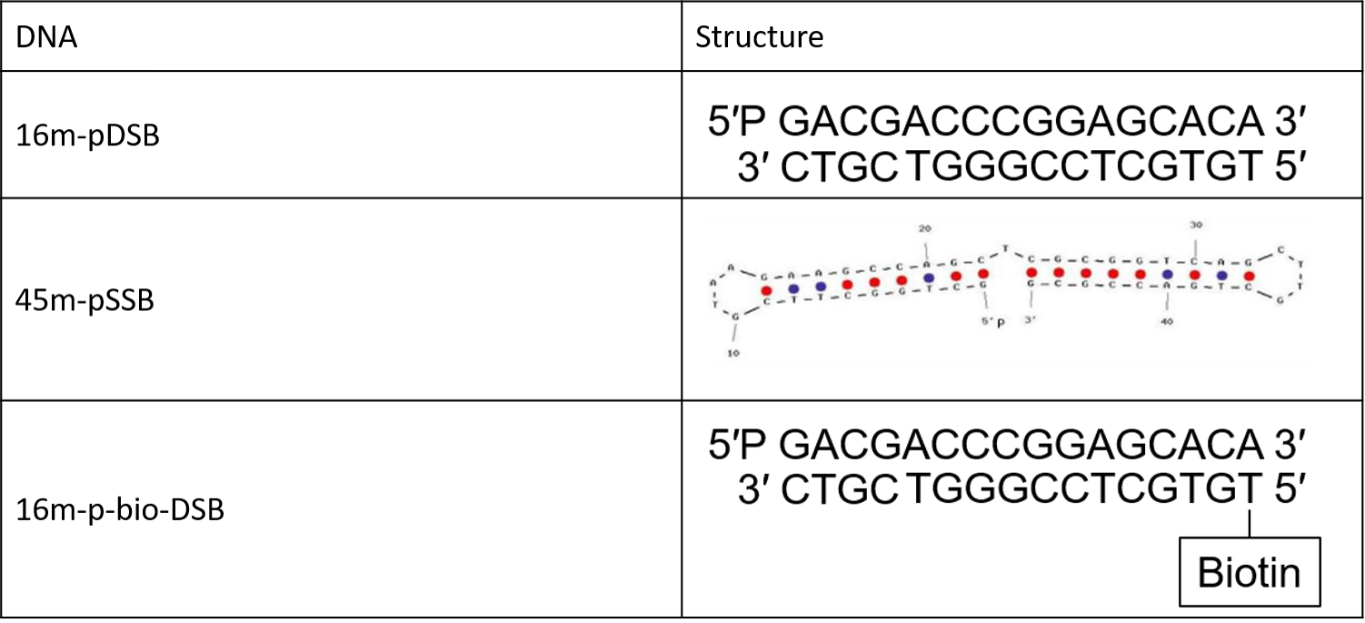

**Supplementary Table 1.** List of DNAs used in the study.
